# Orthohantavirus Diversity in Central-East Argentina: Insights from Complete Genomic Sequencing on Phylogenetics, Geographic patterns and Transmission scenarios

**DOI:** 10.1101/2024.03.25.586579

**Authors:** Daniel Oscar Alonso, Sebastian Kehl, Rocio Coelho, Natalia Periolo, Tomas Poklepovich, Unai Perez-Sautu, Mariano Sanchez-Lockhart, Gustavo F. Palacios, Carla Bellomo, Valeria Martinez

## Abstract

Hantavirus Pulmonary Syndrome (HPS), characterized by its high fatality rate, poses a significant public health concern in the Americas. Phylogenetic relationships of orthohantaviruses in the country were inferred from partial genomic sequences. The objectives of this work were to report new complete viral genomes of the known viral variants associated with HPS cases in Central-East region of Argentina, to assess viral diversity, phylogenetic relationships, and to elucidate the geographic patterns of distribution of each variant. To accomplish this, a detailed analysis was conducted of the geographic distribution of reported cases within the most impacted province of the region. The phylogenetic analysis defined clearly separated clades in the country according to their geographic origin. Andes virus (ANDV) segregated from the rest of the sequences, and those representative from the Central East region, Buenos Aires (BAV) and Lechiguanas virus (LECV), were grouped in the same cluster but segregated in two different branches.

**Importance:** In Argentina, most of the HPS cases were associated with ANDV and closely related viruses distributed in four endemic regions. This work focused on obtaining and studying the complete genome of the orthohantaviruses present in the Central East (CE) region (BAV and LECV). Both viruses were responsible for major and minor outbreaks of person-to-person transmission in the country, and the findings may pave the way to study the impact of genetic determinants of viral transmission and to consider the reclassification of the species *Orthohantavirus andesense*.

## Introduction

Hantavirus pulmonary syndrome (HPS) is a severe zoonotic disease endemic in The Americas, where it shows low incidence but high lethality. Many New World Hantaviruses (NWH) have been described and associated with the disease in all the continent^1^. It is mainly associated with environmental exposure to rodents in rural and wild settings. The infection occurs by inhalation of contaminated aerosols generated by infected rodents that act as reservoirs in nature. Hantaviruses are enveloped, single-strand RNA viruses with tripartite genome consisting of small (S), medium (M), and large (L) segments^2^. Pathogenic Hantaviruses are currently grouped under the genus *Orthohantavirus, family Hantaviridae*. In South America, only five species of orthohantavirus have been recognized by the International Committee on Taxonomy of Viruses (ICTV) despite the fact that 25 distinct viruses were described, most of which have partial genetic information ^3^.

Andes virus (ANDV) was the first orthohantavirus identified as an etiologic agent of HPS in Argentina ^4^. It was associated with up to 50% case fatality rate and person-to-person transmission outbreaks ^5–13^ and considered a global threat to public health. After the description of ANDV, many orthohantavirus were identified in other parts of the country ^14–17^. Several of them were considered as different viruses based on partial genetic information or because they were identified from a different host species; however, the classification of rodent species in the genus *Oligoryzomys*, is still controversial. As several orthohantaviruses identified in Argentina are closely related to ANDV, hereafter referred to as AND-like orthohantaviruses, there is a need to understand the genetic relatedness among them to gain insight into their biological properties. ANDV and AND-like orthohantaviruses were classified under the species *Orthohantavirus andesense*.

ANDV is restricted to Southwestern Argentina and Chile^18^, while AND-like hantaviruses were characterized from CE, northwest and northeast regions of Argentina and surrounding countries^16,19–21^. Given the difficulty to be isolated, the classification of hantavirids was mostly based on genetic relatedness in partial genomic fragments ^22^. The use of partial, non-overlapping fragments could lead to the misidentification of new viruses. Until now, complete genomes were obtained only for ANDV. The absence of L-segment information in hantaviral taxonomic analyses is problematic because it is by far the longest protein encoded by Hantaviruses ^22^. For AND-like Hantaviruses, only few S- and M-segments are available. The study of viral genetic variability and the genetic relatedness among members of the species is then inconclusive. Among HPS cases the most prevalent are Oran virus, BAV and LECV. Regarding remarkable biological properties, BAV is particularly of great concern due to its implication in several outbreaks and suspicion of person-to-person transmission ^23–25^.

The objective of this work was to evaluate the viral divergence in the CE region of Argentina and reconstruct the phylogenetic relationships using complete genomic sequences. For this, the aim was to obtain complete genomes from clinical samples of HPS cases reported in the CE region, obtaining nine high quality, complete viral sequences.

## Results

The CE endemic region of HPS comprises parts of three provinces. The number of reported cases in the region during the period 1996-2022 was 934. Buenos Aires province (BAP) was the most affected according to the number of accumulated cases (n=678, 72.6%). The distribution of 528 cases was studied within localities. The distribution of cases in the province was wide, with highest records in localities placed near riverside areas with shores of the La Plata and Paraná rivers, and other minor rivers that flow into the Atlantic Ocean (Figure 1 A). From 135 localities, 117 (86.7%) reported at least one HPS case, where the number of cases per locality varied from 1 to 134. The most affected were rural areas around La Plata and surrounding localities. Among the cases, 98 were selected for virus characterization, 66.3% were associated with BAV, 26.5% with LEV and 7.1% with PLAV. The pattern of geographic distribution of each virus was different (Figure 1 B). LECV was more frequently found in the northern border of Buenos Aires city along La Plata River through Paraná River, while BAV was widely distributed in the rest of the province from the Delta of La Plata River to the south and southwest. PLAV was found sporadically. The three variants were found cocirculating in the surrounding localities of La Plata (Figure 1 B).

**Figure 1:**
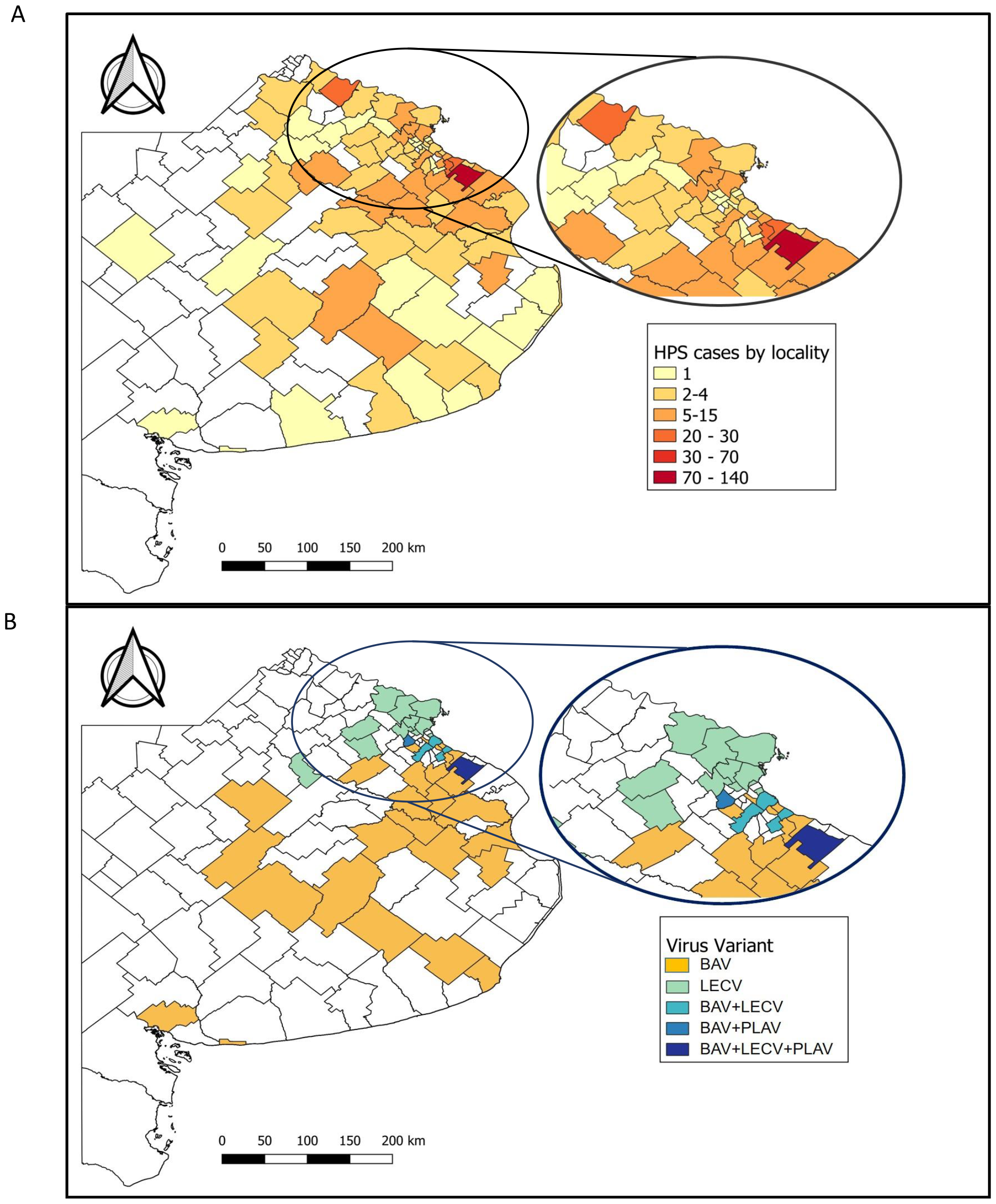
Geographic distribution of Hantavirus Pulmonary Syndrome in Buenos Aires province, Argentina. A: distribution of HPS cases reported by localities in the province during the period 1995-2022 (n=528). B: distribution of viral variants by localities (n=98).

Nine complete (S-, M- and L- segments) and eight incomplete genomes (complete S- and/or M-segments) were obtained. Additionally, an almost complete sequence with only 60.4% of coverage in the L-segment was obtained. Phylogenetic analysis was performed together with available complete sequences in GenBank identified in Argentina, including ANDV and other NWH (Figure 2). The analysis of M- and L- segments showed two main clusters which clearly segregate ANDV-distributed only in southwestern Argentina- (Cluster 1) from AND-like from CE region (Cluster 2) and from the northwest region (Oran virus, only in the M-segment tree). Particularly, in the tree of the S-segment, as there are more complete sequences available, the phylogenetic reconstruction revealed two branches inside Cluster 2 represented by BAV and LECV, with Neembucú, Bermejo and Plata grouped together with LECV (LEC-like variants). In the same tree, other pathogenic viruses were clearly separated in well-defined clusters as the pathogenic Orán and Juquitiba (Clusters 3 and 5, respectively) and the non-pathogenic Pergamino and Maciel (Cluster 4).

**Figure 2:**
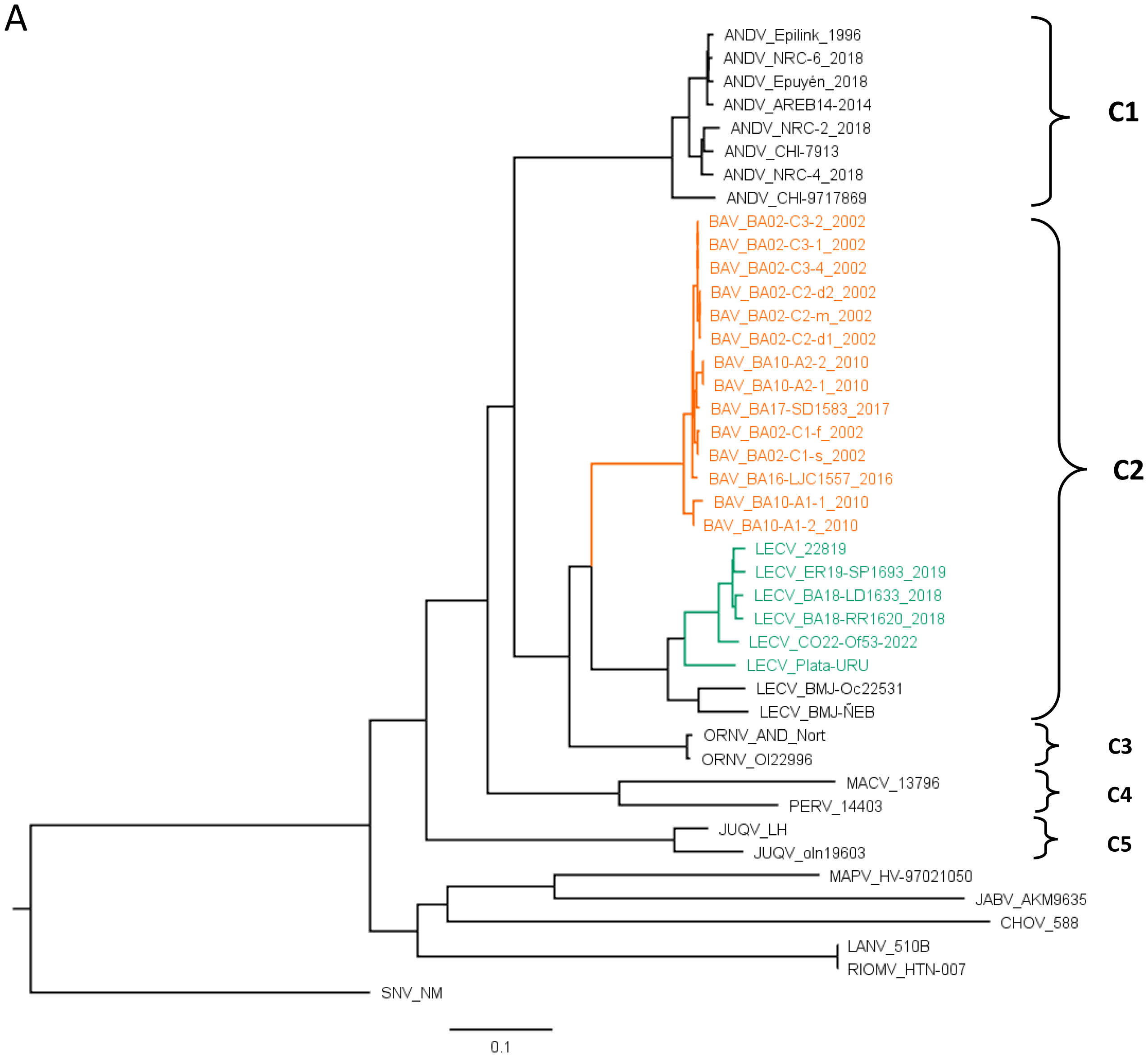
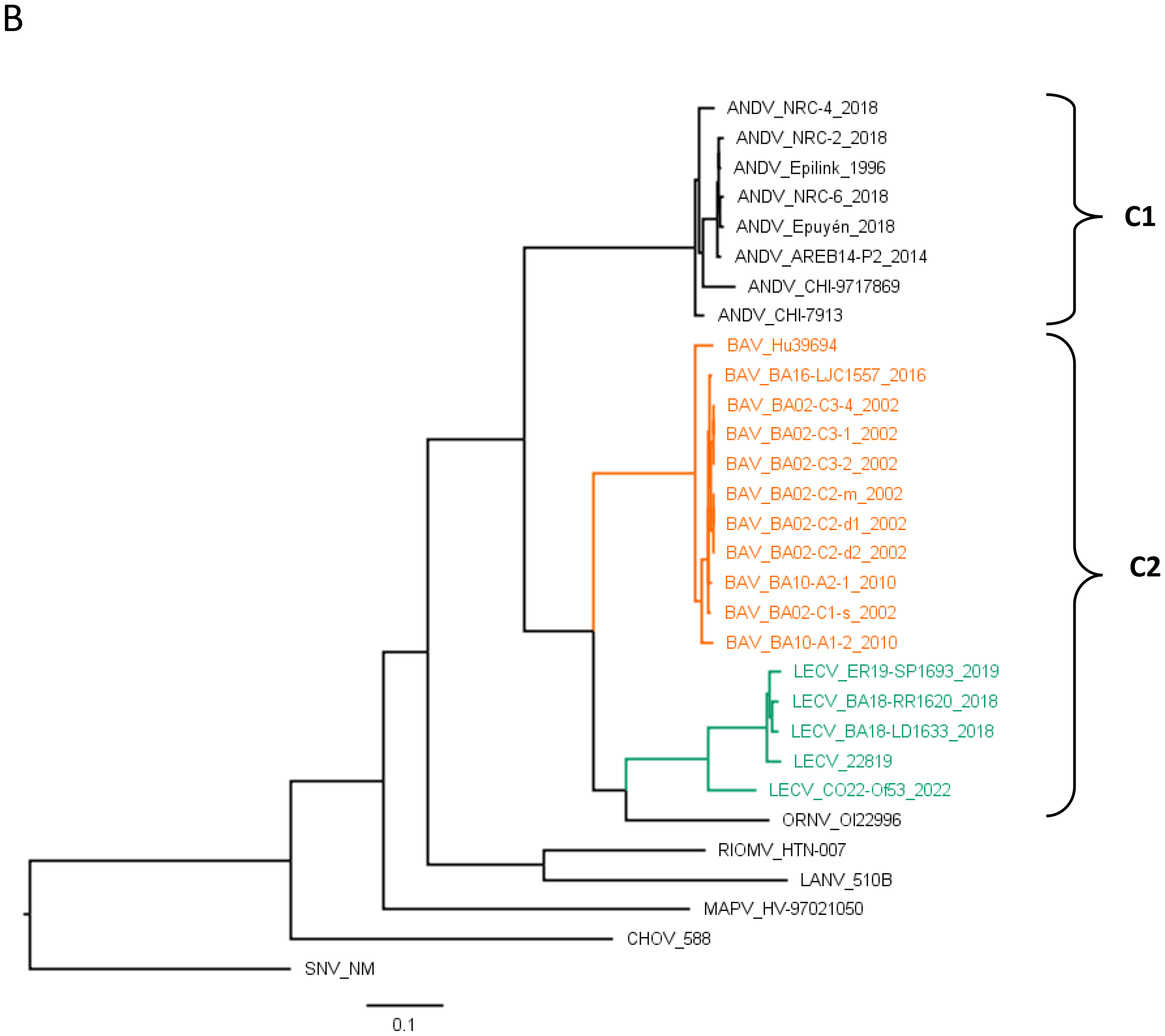
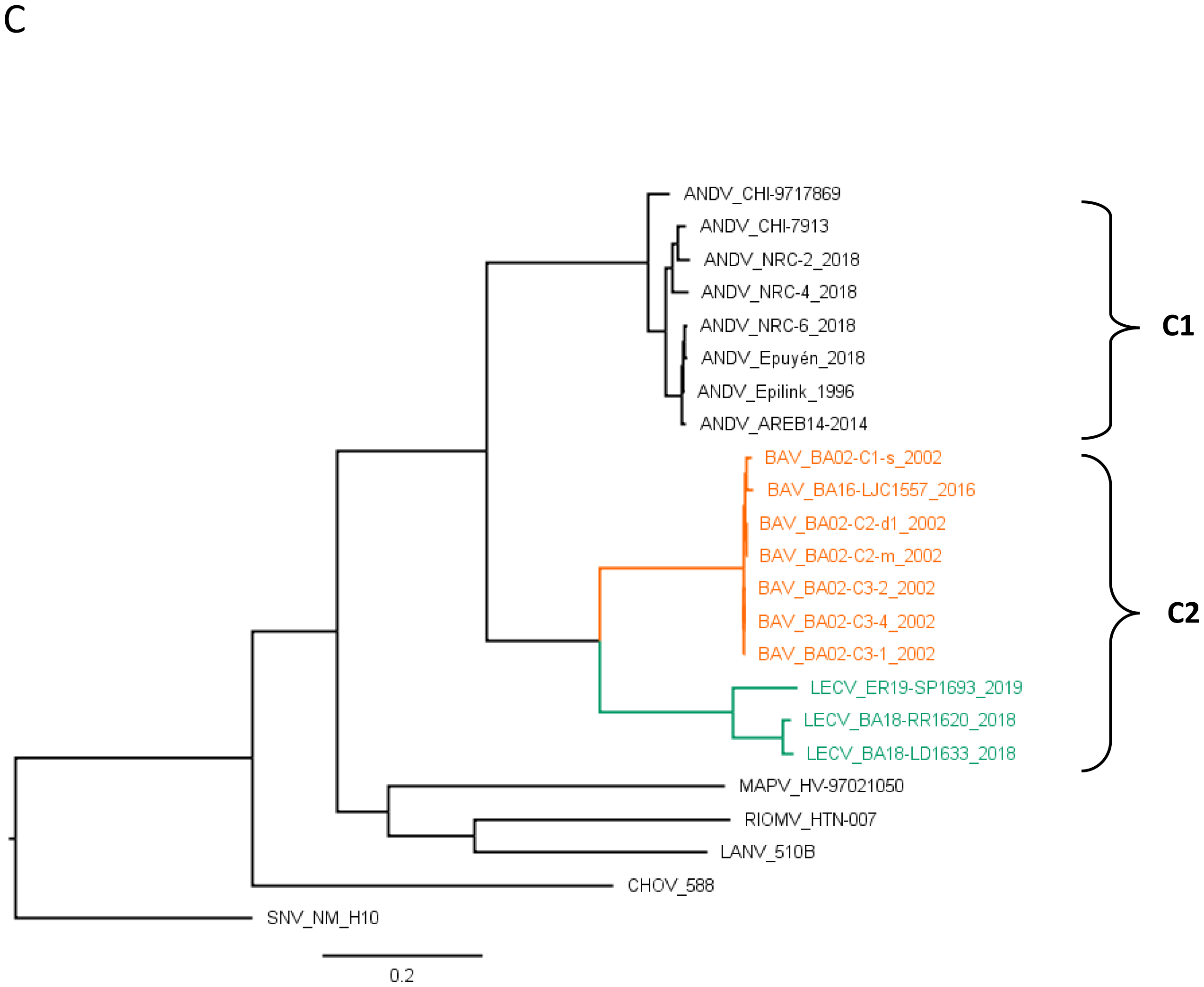
Maximum Likelihood Phylogenetic Analysis based on complete genomes of Orthohantaviruses in Argentina. A: S-segment; B: M-segment; and C: L-segment. The phylogenetic trees were constructed using IQTREE with ModelFinder for model selection, and ultrafast bootstrap analysis with 1000 replicates. The best-fit model according to BIC was GTR+F+I+G4, and this model was used for tree reconstruction. The trees represents the consensus of 1000 bootstrap trees, illustrating the evolutionary relationships among orthohantaviruses.

For viral diversity analysis in our dataset we first estimated the overall genetic variability with the complete genomes (S-, M- and L-segments) of each branch of Cluster 2. The nucleotide divergence range was 0.6-1.2% (n=4) for BAV and 1.8-6.7% (n=3) for LECV. Then, we estimated the divergence between representative viruses of each phylogenetic cluster or branch (Table 2). Compared with ANDV, BAV and LECV diverged 20 and 20.7% respectively, while BAV compared with LECV showed 17.7% at nucleotide level. Considering the three segments separately the divergence at the nucleotide level was similar between them. However, at the amino acid level the divergence was higher in the M segment: ANDV vs. BAV: 8%; ANDV vs LECV: 7%; and ANDV vs. Orán virus: 6%. The divergence in amino acid was remarkably lower for the S segment, indicating a high degree of conservation among all variants present in the country. On the contrary, the S segment non-coding region was the most divergent part of the genome among the different viruses mainly due to specific patterns of insertions and deletions (Table 4).

In previous works, epidemiologically linked HPS cases reported in BAP were analyzed, all of them associated with BAV. Despite 100% of nucleotide identity in partial fragments (total=1000pb from S- and M-segments) between cases in three clusters, person-to-person transmission was confirmed only in one (cluster 1: C1) based on the epidemiological data ^23^. In the present work, a deeper comparative analysis was performed with complete genomes of some of these clustered cases and with complete S-segment sequences of other cases (Table 1) ^24^. In C1, the complete genome of C1-s was obtained, but only the S-segment of C1-f, and therefore only the segment with 100% of nucleotide identity could be compared. Comparing S-segments in other two clusters, A1 and A2, obtained the same result for which the epidemiological investigation could not determine the place of exposure for the secondary cases, as in C1 (Table 3). On the contrary, the intracluster comparison in C2 and C3 revealed changes in the whole genome. In C2, a family cluster in which the symptom onset for all the cases was within a period of 11 days, the three cases analyzed showed at least one change in each segment. The most divergent was C2-d1, which differed from C2-d2 in ten residues. Considering that the coverage of the L-segment of C2-d1 was 60.4%, the total number of changes might be higher. C3, represents another scenario of possible co-exposition in the same house with a maximum period of symptom onset of 20 days between the first and the last cases (C3-1 and C3-4). In this cluster, the differences were up to five residues between C3-2 and C3-4 (Table 3). Intercluster comparisons were performed including non-related cases from the same locality (La Plata) but reported more than 10 years after. The clusters A1 and A2 include geographically and temporally distant cases. The comparisons showed a clear relation between genetic divergence and geographic distance (Table 3).

**Table 1:** Cases of Hantavirus Pulmonary Syndrome selected for sequencing.

| ID | Cluster | Case | Age | Date of onset | Residence | Probable place of infection or risk activity | Sequence coverage |
| --- | --- | --- | --- | --- | --- | --- | --- |
| 1 | C1 | C1-f | 41 | 7/10/2002 | CABA, urban area | Farm in LP, rural area | S complete |
| 2 |  | C1-s | 14 | 8/8/2002 | CABA, urban area | Shared the weekend at his father's house (13 to 14th July/02) | S, M & L complete |
| 3 | C2 | C2-d1 | 12 | 7/24/2002 | LP-El Peligro, rural area (BAP) | LP-El Peligro, rural area (BAP) | S & M complete; L 60.4% |
| 4 |  | C2-d2 | 11 | 7/28/2002 |  |  | S, M & L complete |
| 5 |  | C2-s | NA | NA |  |  | NA |
| 6 |  | C2-m | 40 | 8/4/2002 |  |  | S, M & L complete |
| 7 | C3 | C3-1 | 28 | 8/26/2002 | LP-Abasto, rural area (BAP). | LP-Abasto, rural area (BAP). | S, M & L complete |
| 8 |  | C3-2 | 27 | 8/29/2002 |  |  | S, M & L complete |
| 9 |  | C3-3 | 21 | 9/10/2002 |  |  | NA |
| 10 |  | C3-4 | 30 | 9/14/2002 |  |  | S, M & L complete |
| 11 | A1 | A1-1 | 54 | 3/4/2010 | CABA, urban area | CA (BAP) | S complete |
| 12 |  | A1-2 | 52 | 3/30/2010 | SAP, rural area (BAP) | Contact with A2-1 (husband) | S & M complete |
| 13 | A2 | A2-1 | 58 | 9/5/2011 | BZ, urban area (BAP) | FV, suburban area (BAP) | S complete; M 82% |
| 14 |  | A2-2 | 67 | 9/29/2011 | BZ, urban area (BAP) | Contact with A2-1 (husband) | S complete |
| 15 |  | LJC1557 | 29 | 12/5/2016 | LP-Abasto, rural area (BAP). | LP-Abasto, rural area (BAP). | S, M & L complete |
| 16 |  | SD1583 | 30 | 09/30/2017 | LP-El Peligro, rural area (BAP). | LP-El Peligro, rural area (BAP). | S complete; M 91.2% |
| 17 |  | SP1693 | 37 | 12/24/2018 | Gualeguay, rural area (ERP) | Gualeguay, rural worker | S, M & L complete |
| 18 |  | LD1633 | 49 | 3/6/2018 | CABA, urban area | Alberti (BAP), fishing & camping | S, M & L complete |
| 19 |  | RR1620 | 38 | 1/16/2018 | Don Torcuato (BAP) | Unknown | S, M & L complete |
| 20 |  | URU_1 | NA | 1997 | Uy | Unknown | S complete |

BZ: Berazategui; BAP: Buenos Aires province; CABA = Buenos Aires city; CA: Costa Atlantica; ERP: Entre Rios province; FV: Florencio Varela; LP = La Plata; SAP: San Antonio de Padua; Uy: Uruguay. NA= not available.

**Table 2:**
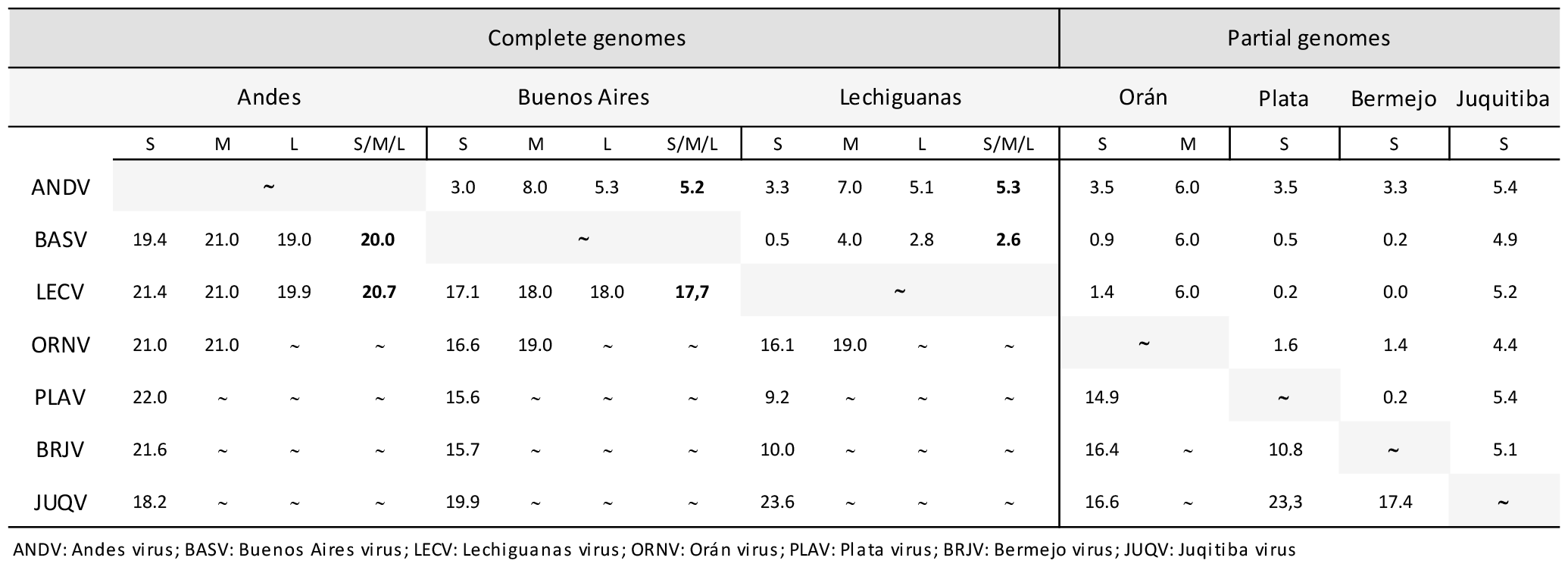
Nucleotide and amino acid comparison between pathogenic orthohantavirus from Argentina.

**Table 3:**
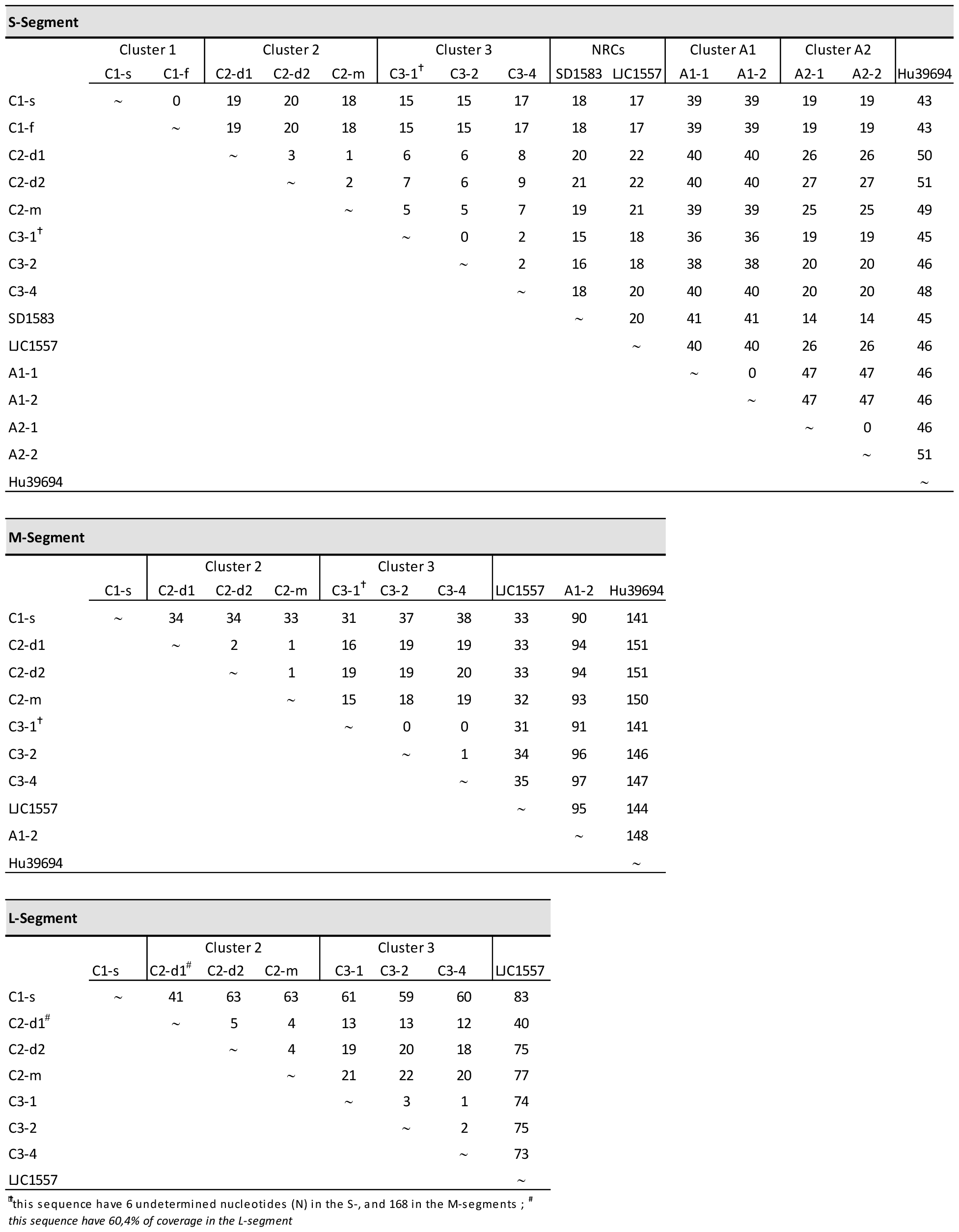
Nucleotide changes between hantavirus Pulmonary Syndrome cases intra and inter-clusters.

**Table 4:**
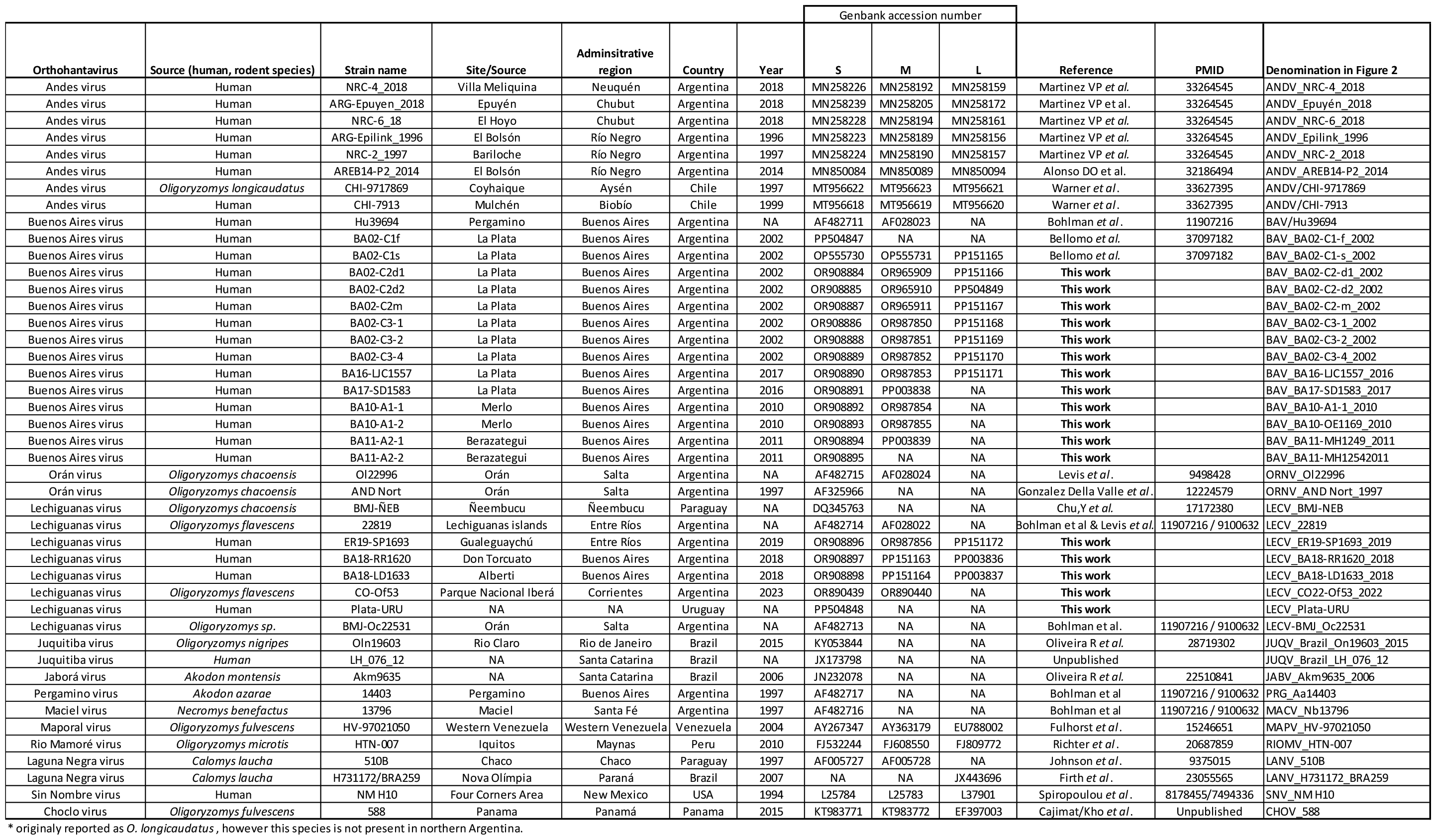
List of sequences and strains utilized in the comparative genomics study of American orthohantaviruses.

## Discussion

Since the identification of the first etiologic agent of HPS in Argentina in 1996, enormous efforts have been channelled to understand viral diversity, routes of transmission and pathogenicity ^5,10,11,23,26,27^. Most of the HPS agents in Argentina and in South America were classified under the species *Orthohantavirus andesense*. Their close relationship with ANDV is a cause for concern in the region because of its ability to spread from person-to-person and high lethality. A large number of incomplete sequences has confounded the understanding of Hantavirus diversity. In this work, for the first time the complete genome of the most prevalent HPS agents in the CE region of Argentina-BAV and LECV-were reported. Nine new complete genomes were obtained from clinical samples of HPS cases that were useful to evaluate genetic variability of each virus. Up to now, only L-segment sequences of ANDV were available. The ICTV Hantaviridae Study Group decided to reassess the entire Hantaviridae family using a stringent criterion which implies to only assess only viruses for which there is S + M + L coding-complete or near-complete sequence information, and this would result in the abolishment of six orthohantavirus species and the declassification (removal from established species) of an additional 11 orthohantaviruses, including LECV and ORNV ^22^.The information provided in this work will help to consider the re-classification of LECV, and the inclusion of BAV as named viruses within *Orthohantavirus andesense*.

The phylogenetic analysis together with complete genomes from Argentina showed two main clusters with all the sequences of ANDV grouped in Cluster 1 and those from the CE region in Cluster 2. Inside Cluster 2, BAV and LECV are defined in two branches. BAV and LECV differed from ANDV only in 5.2 and 5.3% in amino acids. The phylogenetic reconstruction of the S-segment showed a similar topology for the other segments, but as there are a higher number of complete genomes, it allows the distinction of a third cluster, represented by Juquitiba, a virus identified in the northeast region of Argentina and Brazil ^16,28,29^. Other sequences as Bermejo, Plata and Neembucú, which differed between them in up to 11% in nucleotides, grouped in the subcluster together with LECV. Then, all these variants which showed a minimal divergence at the amino acid level should be considered strains of LECV. However, for the definite classification of these strains, complete sequences should be obtained.

It was estimated that almost 70% of HPS cases in BAP were caused by BAV, which has a wider geographic distribution than LECV. Nonetheless, while BAV is restricted to BAP, LECV showed a wider distribution outside BAP to the north (even in the northeast region of Argentina) and to the east (Uruguay) ^16,30,31^. Despite the distribution showed in this work, two cases of BAV were previously reported in the northwest region, evidencing the need to address viral genotyping studies in the whole country ^32,33^. The distinctive geographic distribution pattern of BAV and LECV are probable indicators of favourable ecological conditions of different reservoir hosts, however, the rodent host of BAV has never been identified. Therefore, further efforts should be conducted in order to identify this reservoir.

Despite that BAV and LECV showed similar levels of divergence from ANDV, only BAV was implicated in person-to-person transmission and in several clustered cases as well ^6,23–25,34^. In previous works 100% nucleotide identity was found in partial fragments of viral genomes (around 10% of the genome) in three clusters of epidemiologically linked cases. Person-to-person transmission in paired cases was postulated based on accurate epidemiologic information that probed absence of exposure to the same rodent population. In this work, 100% of nucleotide identity we found in the S-segment in three clusters adding genetic evidence in favour of person-to-person transmission. In contrast, in two clusters where co-exposure was evident, the whole genome was analyzed and several nucleotide changes were found. In C2, the short period of symptom onset between all the members of the family and the number of changes found, strongly suggest exposure to different infected rodents of the same population. In C3, the infections could have occurred from different or the same rodent, but at least one of the cases could have been infected from person-to-person, however a chain of transmission could be ruled out due to the absence of clear patterns of changes between cases. This contrasted with our findings in the sustained ANDV person-to-person outbreak that occurred in Epuyén in 2018, in which up to two mutations appeared in only six patients in the chain of transmissions that involved 33 cases^10^. A question that frequently arises from genomic analysis when facing clusters of cases is which would be the threshold of changes to differentiate person-to-person transmission from co-exposure to the same infected rodent. The answer remains elusive and requires deeper studies involving rodent reservoir populations. Nevertheless, the findings reported here are important and could help to resolve uncertainties in future outbreaks.

In conclusion, high quality and complete genomic sequences were obtained of many isolates of two viruses responsible for the majority of the HPS cases in the CE region of Argentina. Our results showed that both viruses diverge from each other in 17.7% and 2.6% in nucleotide and amino acid levels respectively, have different geographical distribution patterns and also differ in the biological property to spread person-to-person, only described for BAP to date. Further efforts should be focused on obtaining new complete genomes from cases and rodent host populations to fill the gaps in Hantavirus classification to understand viral diversity and biological traits such as host range, routes of transmission and pathogenesis. Finally, complete genomic analysis has become a critical tool for the distinction of viral spillover from person-to-person transmissions. This study enhances our understanding of the genetic diversity, geographical spread, and transmission dynamics of orthohantaviruses involved with HPS in Central Argentina.

## Materials and Methods

Samples and clinical/epidemiological information are available in the laboratory, as National Reference Laboratory for Hantavirus. A retrospective and transversal study of case distribution from 1995 to 2022 was performed in the CE region including all the cases reported in the region (https://sisa.msal.gov.ar/sisa/). In particular, Buenos Aires, the most affected province of the CE region, was selected to construct a map showing the geographical distribution of HPS cases and viral variants. Buenos Aires province is located between 33°40′35′′ and 41°8′49′′ S and between 56°24′42′′ and 63°10′35′′ W, covering a total surface of 307,571 km2 ^35^. It is divided into 135 localities. Confirmed cases with records of residence as the most probable place of infection and without recent history travel were selected (n=528). The distribution of cases was mapped using QGIS software (3.16). For the construction of the map of variants distribution, viral genomes previously characterized from HPS cases (n=98) were used. The genetic characterization was performed on partial fragments as previously described ^32^.

In order to obtain complete genomes of all the variants circulating in the area, 18 cases were selected (Table 1), including three characterized as LECV, one Plata virus and 14 BAV; most of the selected BAV cases were epidemiologically related and were involved in five clusters reported previously^23,24^. For the genetic analysis, total RNA was extracted from blood samples using *in-house* methods based on manual extraction with TRIzol™ Reagent. For whole viral genome sequencing, libraries were prepared by two different strategies, target-capture enrichment as previously described ^10^ and an amplicons based method using a two-step RT-PCR to enrich vRNA which could be followed by a second round of PCR if needed, and the subsequent library preparation. Pooled libraries were sequenced on the Illumina MiSeq, NextSeq or NovaSeq sequencing platforms (Illumina, San Diego, CA). Bioinformatic analysis on fastQ resulting files were performed as previously described Martinez et al 2020 supplementary material. Only bases with a Phred quality score >Q30 and a minimum of 10X coverage were used for consensus calling. Consensus genome sequences from cases were aligned using MAFFT v.7.397.

The diversity of the viral population was then estimated according to two parameters: the total number of individual nucleotide changes in each genomic sequence and the total number of amino acid changes in the coding regions of each segment. The percentage of divergence was determined by alignment analysis with the basic local alignment search tool, BLAST ^36^. BlastN was selected for the comparison of more dissimilar sequences. The remaining parameters were set by default.

